# Mating-type locus structure affects gene expression in unidirectional mating-type switching fungi

**DOI:** 10.1101/2025.10.10.681704

**Authors:** Frances A. Lane, Brenda D. Wingfield, Michael J. Wingfield, P. Markus Wilken

## Abstract

Fungal species are typically either fully self-fertile or self-sterile, but some filamentous ascomycetes can commonly transition from self-fertility to self-sterility through unidirectional mating-type switching. In these fungi, the structure of the mating-type (*MAT1*) locus governs sexual behaviour: MAT-2 self-fertile individuals retain both *MAT1-1* and *MAT1-2* genes, while MAT-1 self-sterile isolates lose *MAT1-2* genes during switching. A third type of isolate morphology also occurs under laboratory conditions: these are self-sterile isolates which retain both *MAT1-1* and *MAT1-2*, but are unable to switch mating type. These are commonly referred to as MAT-2 self-sterile isolates. Two of the mating-type (*MAT*) genes, one of which is deleted during switching, encode transcription factors known to regulate not only the sexual cycle but also genes unrelated to mating. To test how *MAT1* structural variations affects gene expression, we studied *Ceratocystis albifundus*, a species that switches mating type. To minimise variability caused by intraspecific genetic differences, two self-sterile isolates (MAT-1 and MAT-2 self-steriles) were derived from the same MAT-2 self-fertile parent, making all three isolates genetically identical except at the *MAT1* locus. Comparative transcriptomic analyses revealed that the MAT-2 self-fertile, MAT-1 self-sterile and MAT-2 self-sterile isolates all exhibited distinct expression patterns, including differences in *MAT* genes, the pheromone–receptor pathway, and other genes not directly linked to mating. The results show that *MAT1* locus structure influences gene expression more broadly than those only related to the sexual cycle.

## Introduction

Sexual reproduction in ascomycetes is controlled by a single genomic region known as the mating-type (*MAT1*) locus (Wilson et al. 2021). The gene content and overall architecture of the *MAT1* locus drives mating behaviour and reproductive strategy. Is its simplest form, the *MAT1* locus is defined by the *MAT1-1-1* and/or *MAT1-2-1* genes (Turgeon and Yoder 2000; Wilken et al. 2017), with both encoding transcription factors that influence gene expression, ultimately enabling the sexual cycle (Dyer et al. 2016; Herskowitz 1989; Wilson et al. 2021). The gene content and structure of the *MAT1* locus can vary across fungi, and in some cases even shows variation within members of a single species (Butler 2007). This variation accounts for most of the diversity of fungal mating strategies.

Most fungi have either a heterothallic or homothallic mating strategy (Bennett and Turgeon 2016; Butler 2007; Debuchy et al. 2010). Heterothallic species are self-sterile as they require a partner of opposite mating type to complete the sexual cycle. In contrast, homothallic fungi can self-fertilize to produce sexual spores, but this ability can arise through several distinct genetic mechanisms (Dodge 1957; Glass and Smith 1994; Ni et al. 2011; Perkins 1987; Wilson et al. 2015). The most well known of these is primary homothallism where all the genes driving mating type are present in a single isolate (Pöggeler et al. 1997). Other mechanisms also enable self-fertility in fungi, such as mating-type switching, where genetic alterations in the *MAT1* locus drives self-fertility. While bidirectional mating-type switching is well characterised in yeasts (Haber 2012; Hanson and Wolfe 2017), unidirectional switching in filamentous ascomycetes is comparatively less well understood.

Unidirectional mating-type switching produces both self-fertile and self-sterile progeny from a single sexual event, whether by selfing or outcrossing (Perkins 1987). This behaviour has only been described in a small number of species residing in four fungal families; *Ceratocystidaceae* (Harrington and McNew 1997), *Glomerellaceae* (Wheeler 1950), *Hypocreaceae* (Mathieson 1952), and *Sclerotiniaceae* (Uhm and Fujii 1983a; 1983b). In two of these species, *Chromocrea spinulosa* (*Trichoderma spinulosum*) and *Sclerotinia trifoliorum*, the ascospores that give rise to self-fertile and self-sterile strains differ in size (Uhm and Fujii 1983a; 1983b; Wheeler 1950). While this spore dimorphism is not known in any of the species that undergo unidirectional mating-type switching in the *Ceratocystidaceae* (Harrington and McNew 1997), there have been some reports that the self-fertile isolates are more pathogenic (Lee et al. 2015). This is not surprising as several studies on fungi residing in other families have shown that mating-type genes directly influence many other aspects of their biology, including vegetative growth rate, conidial structure and pathogenicity (Lockhart et al. 2005; Yong et al. 2020; Zhan et al. 2007).

Recent work has highlighted key genetic mechanisms underlying unidirectional mating-type switching in fungi. Self-fertile isolates appear to exist as heterokaryons that differ only at the *MAT1* locus (Yun et al. 2017). One version of the *MAT1* locus contains both *MAT1-1* and *MAT1-2* genes, with the *MAT1-2* genes typically inserted within the *MAT1-1-1* gene, disrupting its function (Wilken et al. 2024; Xu et al. 2016; Yun et al. 2017). The alternative version contains only *MAT1-1* genes and, when present as a homokaryon, is self-sterile. Unidirectional mating-type switching produces this self-sterile form by deleting the *MAT1-2* genes and one of the flanking direct repeats (Wilken et al. 2014; Witthuhn et al. 2000), thereby restoring *MAT1-1-1* function (Wilken et al. 2024). Despite their self-sterility, these isolates retain the ability to mate and can outcross with self-fertile strains to produce viable offspring (Harrington and McNew 1997; Yun et al. 2017). These findings highlight the central role of unidirectional mating-type switching in generating both self-fertile and self-sterile isolates, ultimately enabling sexual reproduction to occur.

Interestingly, some species in the *Ceratocystidaceae*, including members of the genus *Ceratocystis*, have an unusual self-sterile morphotype (Harrington and McNew 1997). In this case, individuals are referred to as MAT-2 self-sterile isolates and have exclusively been described in laboratory strains. PCR analysis has shown that these isolates are homokaryotic, carrying a *MAT1* locus with both *MAT1-1* and *MAT1-2* genes similar to the MAT-2 self-fertile isolates (Engelbrecht and Harrington 2017; Oliveira et al. 2015; Wilken 2015). Although the cause of sterility in MAT-2 self-sterile mutants is unknown, they have been used in mating and hybridisation studies where they exclusively act as a male mating partner (Fourie et al. 2019; Johnson et al. 2005; Oliveira et al. 2015).

*Ceratocystis albifundus* is emerging as a genetic model for studies on members of the genus *Ceratocystis*. Current research efforts for this species include the development of an *Agrobacterium*-mediated transformation system, a population-level genomic study, and the identification of its pheromone-receptor system (Danki et al. 2024; Lane et al. 2025; Sayari et al. 2019; van der Nest et al. 2019). The MAT-2 self-fertile, naturally occurring MAT-1 self-sterile, and laboratory-derived MAT-2 self-sterile isolates have been described (Lee et al. 2018). The *MAT1* locus of MAT-2 self-fertile isolates contains complete copies of *MAT1-1-1*, *MAT1-1-2*, *MAT1-2-1*, and *MAT1-2-7* (Lee et al. 2018; Wilken et al. 2024). During unidirectional switching, the *MAT1-2* genes are deleted, producing a MAT-1 self-sterile isolate. Although *MAT1-1-1* is structurally intact in the self-fertile morphotype, the presence of *MAT1-2* genes appears to separate it from its promoter. Deletion of the *MAT1-2* region joins *MAT1-1-1* to the promoter, enabling its expression. However, the expression profile of the mating-type genes and other genes involved in the sexual cycle of this fungus remains to be explained.

The ability of *Ceratocystis* species to produce both self-fertile and self-sterile versions of the mating-type locus within an otherwise uniform genetic background makes them ideal to study how *MAT1* structure influences gene expression. In these fungi, the phenotype of an isolate, whether it is self-fertile or self-sterile, is determined directly by the arrangement of the *MAT1* locus. The aim of this study was to explore how differences at this single locus influences gene expression in *C. albifundus*. All isolates used were genetically identical except for the *MAT1* region, making it possible to consider how self-sterile isolates, which functionally mimic heterothallic mating types, differ from each other and from a self-fertile isolate that carries both idiomorphs. By examining the transcriptomes of the three isolate types, we gained insights into how *MAT* genes are expressed within different locus structures, and how variation at the *MAT1* locus alone can shape overall genome-wide gene expression.

## Materials and methods

### Annotated genome assembly for functional analysis

The genome assembly of *Ceratocystis albifundus* isolate CMW 4068, a MAT-2 self-fertile isolate, was obtained from NCBI (GenBank accession GCA_002742255) and was annotated using data provided by van der Nest et al. (2019). The *MAT1*, pheromone and pheromone receptor loci were screened to confirm that the annotations included all genes expected in these regions (Lane et al. 2025; Wilken et al. 2024), and, where missing, they were manually added. Annotations for the direct repeats were also added to the *MAT1* locus.

The mating-type locus from the genome was edited manually to produce two loci that were representative of the transcriptionally active *MAT* gene structure (Wilken et al. 2024). To do this, the *MAT1* locus was located based on a tBLASTn search using the *MAT* genes described in *C. albifundus* (Wilken et al. 2024). This locus was edited by removing the complete *MAT1-2* region, that consisted of the *MAT1-2-1* and *MAT1-2-7* genes and a single direct repeat. This produced the *MAT1* locus corresponding to the MAT-1 self-sterile version with the *MAT1-1-1* gene linked to its promoter (Wilken et al. 2024). To retain the *MAT1-2* genes, the removed region was artificially included in the genome assembly as a freestanding contig. The genome assembly was functionally annotated using the OmicsBox (Bioinformatics Made Easy, BioBam Bioinformatics) Blast2GO suite which included a Diamond BLAST followed by InterProScan, and EggNOG analyses (Blum et al. 2021; Gotz et al. 2008; Huerta-Cepas et al. 2017; Huerta-Cepas et al. 2016).

### Generation of self-sterile cultures

*Ceratocystis albifundus* isolate CMW4068 was obtained from the culture collection (CMW) of the Forestry and Agricultural Biotechnology Institute (FABI) based at the University of Pretoria. This culture was grown at 25°C on 2% malt extract agar (MEA; Biolab, Merck, Johannesburg, South Africa) supplemented with 150 mg/ml thiamine and 100 mg/ml streptomycin (TS; SIGMA, Steinheim, Germany). After approximately two weeks of incubation, ascomata having drops of ascospores at their apices, typical of self-fertility in this species, were observed. This isolate was designated as MAT-2 self-fertile in the study. Single ascospore isolates were generated from this culture using serial dilution as described in Krämer et al. (2021). Single ascospore cultures that did not produce ascomata after two weeks of incubation were treated as putatively self-sterile. These isolates were subcultured for two more rounds by transferring a block of agar covered with mycelium block to a fresh MEA-TS plate. When these cultures remained sterile, they were further screened for the presence of one or both versions of the mating-type locus.

DNA was isolated from all potential self-sterile cultures and the corresponding self-fertile culture using a CTAB protocol (Krämer et al. 2021), and subjected to PCR screening to determine *MAT1* locus structure. Two primer sets targeting the *MAT1* locus were used to distinguish between the MAT-2 self-fertile and MAT-1 self-sterile versions, with two additional gene-specific primer sets included for confirmation (Supp. Table 1; Suppl. Fig. 1). All PCR amplifications were carried out in separate 25 μl reactions consisting of 10 – 50 ng of template DNA, 1 U KAPA Taq DNA Polymerase (KAPA Biosystems, Boston, MA, USA), 1 X KAPA Taq Buffer A, 0.2 mM dNTP mix and 0.4 μM of each primer. The DNA from the MAT-2 self-fertile isolate served as a positive control as these cultures are expected to carry both versions of the *MAT1* locus and all *MAT* genes (Wilken et al. 2024). Amplification reactions were performed as described in Wilken et al. (2024) using an annealing temperature of 55°C. PCR amplicons were visualised via gel electrophoresis, and the results were used to identify the self-sterile isolates as either MAT-1 self-sterile or MAT-2 self-sterile.

### RNA isolation and sequencing

A single isolate of each of the three isolate types (MAT-2 self-fertile, MAT-1 self-sterile and MAT-2 self-sterile) were selected for further analysis. For each isolate type, nine petri dishes containing 2% MEA-TS covered with a sterile cellophane sheet (BioRad, Johannesburg, South Africa) were inoculated and incubated for 13 days in the dark at 25°C. For RNA extraction, the mycelium growing on the surface of the cellophane from three plates was pooled into a single extraction, and this was done in triplicate for the nine cultures per isolate (resulting in three RNA samples for each isolate type). RNA extractions were performed using the RNeasy Plant extraction kit (Qiagen, Limburg, The Netherlands) following the manufacturer’s instructions apart from using the RLC buffer and including a DNase extraction step using a DNase-I kit (RNase-Free DNase Set, Qiagen, Limburg, The Netherlands). RNA sample quantity and quality was assessed using gel electrophoresis and a NanoDrop ND-1000 (ThermoScientific, Waltham, USA). RiboLock RNase Inhibitor (ThermoScientific, Waltham, USA) was added to each sample before storage at -80°C.

RNA library preparation and sequencing was performed by the Agricultural Research Council Biotechnology Platform (ARC-BTP, Onderstepoort, South Africa). The library was produced using an Illumina TruSeq Stranded Total RNA kit (Illumina, San Diego, CA, USA), which includes the removal of rRNA and capturing of both coding and non-coding RNA, the quality of which was assessed using a Qubit 3.0 fluorometer (ThermoFisher Scientific, Waltham, MA, USA) and LabChip GX Touch (Perkin Elmer,Waltham, MA, USA) instrument. The RNA sequencing was completed on an Illumina HiSeqX instrument (Illumina, San Diego, CA, USA).

### Expression analysis

CLC Genomics Workbench (v. 22.0; CLC Bio, Aarhus, Denmark) was used to map the raw reads to the target genome assembly. Raw reads were trimmed using the default settings (Suppl. Table 2) of the built-in “Trim Reads” tool and the quality was assessed using the QC function. RNA reads of each isolate type were mapped to the modified *C. albifundus* annotated genome using the “RNA-Seq Analysis” tool. Default settings were used except for the length fraction which was set to a minimum of 0.5 (Suppl. Table 3).

The R studio package zFPKM (v. 1.26.0) was used to normalise the mapped reads across samples and to identify genes that were expressed relative to background gene expression (Hart et al. 2013). Fragments per kilobase of transcript per million mapped reads (FPKM) values for the coding sequence (CDS) of each gene generated by CLC Genomics Workbench was normalised using zFPKM. The mean zFPKM value across the three replicates for each sample type was calculated and genes with a mean zFPKM of more than or equal to -3 (as recommended by HART *et al*. 2013) were considered expressed.

Differential gene expression was performed by DESeq2 (v. 1.44.0; Love et al. 2014) using filtered CDS datasets of raw counts and comparing the MAT-2 self-fertile to both self-sterile isolates separately as well as comparing the self-sterile isolates to each other. Genes that were not expressed (based on the zFPKM values) in both isolates being compared were removed to reduce background noise. The *MAT1-2-1* and *MAT1-2-7* genes were also removed in comparisons with the MAT-1 self-sterile isolates as these genes are absent in these genomes and thus would not show any expression.

Genes were considered significantly differentially expressed between sample types if there was a log2 fold change (log2 FC) of more than 0.58 or less than -0.58 (≈ 1.5 fold change) with a adjusted p-value of less than 0.05 (Joubert et al. 2024). The adjusted CDS datasets of raw counts as well as the results of DESeq2 were used to produce principal component analyses (PCA) plots and heatmaps (Suppl. File 1). Amongst the top ten genes that showed the highest upregulation from each isolate comparison, any described as “hypothetical” based on BLAST results were translated and conserved domains were identified using a Pfam (Pfam-A database v. 37.2) search in CLC Main to discern possible functions for these gene products. The expression of the mating-type and pheromone/receptor pathway genes were also visualised in a heatmap using rlog-transformed normalised counts with gene clustering using default settings on pheatmap (v. 1.0.12).

Gene Set Enrichment Analysis (GSEA; Mootha et al. 2003; Subramanian et al. 2005) was used to determine the overall pathway enrichment of each comparison. The sign(log_2_FC) x -log_10_(p-value) from the DESeq2 output was used as the ranking metric (Stavely et al. 2023). Gene sets were considered enriched if they had an FDR p-value of less than 0.25, a normalised enrichment score (NES) of more than 1 or less than -1 and a nominal (NOM) p-value of less than 0.05. The top 20 differentially expressed gene sets (for both up and down regulation; based on the NOM p-value) were visualised using ggplot2 (v. 3.5.1). Unique and shared differentially expressed genes were also identified using an online Venn diagram tool (https://bioinformatics.psb.ugent.be/webtools/Venn/) and then gene set enrichment for these groupings was determined using the R clusterProfiler package (v. 4.12.6) using the Benjamini & Hochberg adjustment method and a p-value cutoff of 0.05 (Yu et al. 2012).

The mapping of the reads from the mating-type genes, pheromone and pheromone-receptor genes were further analysed. To do this, the three RNA datasets for each isolate type were individually mapped to the MAT-2 self-fertile and MAT-1 self-sterile versions of the *MAT1* locus. The mapping results were inspected for anomalies in the read distribution across the gene, as well as for possible alternative splicing events.

## Results

### Annotated genome assembly for functional analysis

The genome of the MAT-2 self-fertile isolate CMW 4068 that contained both the MAT-2 self-fertile and MAT-1 self-sterile *MAT1* locus versions, was used to map the RNA reads from all three datasets. After manually annotating the a-pheromone and *MAT1-2-7* genes, a total of 7,024 gene annotations were present in the genome assembly. Of these genes, the Blast2GO pipeline identified 2,185 as hypothetical proteins, with 1,262 of these assigned GO terms. A further 223 genes lacked homology to genes in the searched database and were not assigned a descriptive name or GO term. These genes were subsequently referred to as putative genes in this study.

The mating-type locus was identified on contig MAOA02000032 and was identical to the self-fertile *MAT1* locus version previously described (Wilken et al. 2024). It contained four mating-type genes, *MAT1-1-2*, *MAT1-2-1*, *MAT1-2-7* and *MAT1-1-1*. Two 84 bp direct repeats, flanking the *MAT1-2* genes, were also identified. The first direct repeat was located between the *MAT1-1-2* and *MAT1-2-1* genes while the second formed part of the 5’ end of the *MAT1-1-1* open reading frame.

The region flanked by the direct repeats would have been deleted through mating-type switching, and was 3,348 bp in size. This region included *MAT1-2-1* and *MAT1-2-7*, and a single copy of the repeat. This was removed from the contig and inserted into the genome assembly as a stand-alone contig. By doing so, the self-fertile *MAT1* locus version present in the genome assembly was replaced by the self-sterile version where the *MAT1-1-1* gene is placed directly downstream of its promoter (Wilken et al. 2024). The *MAT1-2* genes were retained in a separate contig together with their immediately flanking regions to avoid disrupting any potential promoter sequences. This allowed for more accurate mapping of the expression data.

### Generation of self-sterile cultures

The isolate of *C. albifundus* (CMW 4068) produced abundant fertile ascomata when cultured in isolation, and was confirmed as a MAT-2 self-fertile individual. Forty single-spore colonies were derived from this isolate, of which five consistently lacked ascomata and were treated as putatively self-sterile. PCR screening showed that one isolate possessed only the MAT-1 self-sterile version of the *MAT1* locus, with no amplicons matching the MAT-2 self-fertile locus or *MAT1-2-1* gene being present. This isolate was subsequently treated as a MAT-1 self-sterile. The PCR results for the remaining four isolates consistently produced the expected amplicons for the *MAT1-1-2* and *MAT1-2-1* genes, as well as the MAT-2 self-fertile *MAT1* locus version. The structure of the *MAT1* locus, combined with the self-sterility was indicative of a MAT-2 self-sterile isolate. One of these was randomly chosen for the remainder of the study. The MAT-1 and MAT-2 self-sterile isolates have been deposited into the culture collection (CMW) of the Forestry and Agricultural Biotechnology Institute (FABI) at the University of Pretoria.

### RNA isolation and sequencing

RNA of good quality and high concentration was extracted from the MAT-2 self-fertile, the MAT-1 self-sterile as well as one of the MAT-2 self-sterile isolates (Suppl. Table 4). RNA sequencing produced an average of 37.1 million raw reads for the MAT-2 self-fertile, 44.7 million reads for the MAT-1 self-sterile, and 35.6 million reads for the MAT-2 self-sterile (deposited in the NCBI Sequence Read Archive). After quality trimming, at least 99.997% of the reads were retained. On average, 88.58% (for the MAT-2 self-fertile), 88.79% (for the MAT-1 self-sterile) and 90.43% (for the MAT-2 self-sterile) of the reads mapped to the genome (in pairs and in broken pairs), representing an average of 10.2 million, 10.0 million and 7.4 million transcripts for each sample, respectively.

### Expression analysis

Of the 7,024 predicted genes in the genome, 6,495 were consistently expressed across all isolates, indicating strong transcriptome coverage and a high level of overall expression. To reduce background signal and enhance differential expression analysis, genes not expressed in both isolates of a given comparison, or absent from the MAT-1 self-sterile genome (i.e., *MAT1-2-1* and *MAT1-2-7*), were removed. This included 339 genes not expressed in any isolate, and a small number of additional genes specific to each comparison: 41 for the self-fertile vs. MAT-1 self-sterile, 12 for the self-fertile vs. MAT-2 self-sterile, and 53 for the MAT-1 vs. MAT-2 self-sterile comparison. The extensive overlap in expressed genes suggested that there was a low risk of failing to detect expression differences.

Filtering for expressed genes and principal component analysis (PCA) confirmed data quality, replicate consistency, and clear separation between isolate types. Genes not expressed across isolate types were removed to reduce technical artifacts, resulting in refined gene sets for each comparison to improve differential expression analysis. The PCAs revealed clustering of technical replicates for each sample with the PC1 accounting for 80 and 83% variance between the two self-sterile isolate types and the MAT-2 self-fertile isolate with which it was compared (Suppl. Fig. 2). While some variation was observed between technical replicates (Suppl. File 2), the average intra-replicate distance was significantly smaller than the inter-isolate distance, supporting the reproducibility of the technical replicates used. Due to the small sample size and the clear distinction observed between isolate types, this variation was deemed unlikely to impact the overall conclusions.

Across all isolates, genes associated with the mating-type and pheromone signalling pathways displayed marked differences in expression levels. Both secondary mating-type genes (*MAT1-1-2* and *MAT1-2-7*) and the pheromone receptor genes were expressed at low levels (Fig. 1). In contrast, *mak2* (involved in signal transduction), as well as *ram2* and *ste24* (involved in a-pheromone post-translational modification), showed consistently high expression relative to the other mating-related genes. Notably, neither the a-nor α-pheromone genes were expressed in any isolate.

**Figure 1:**
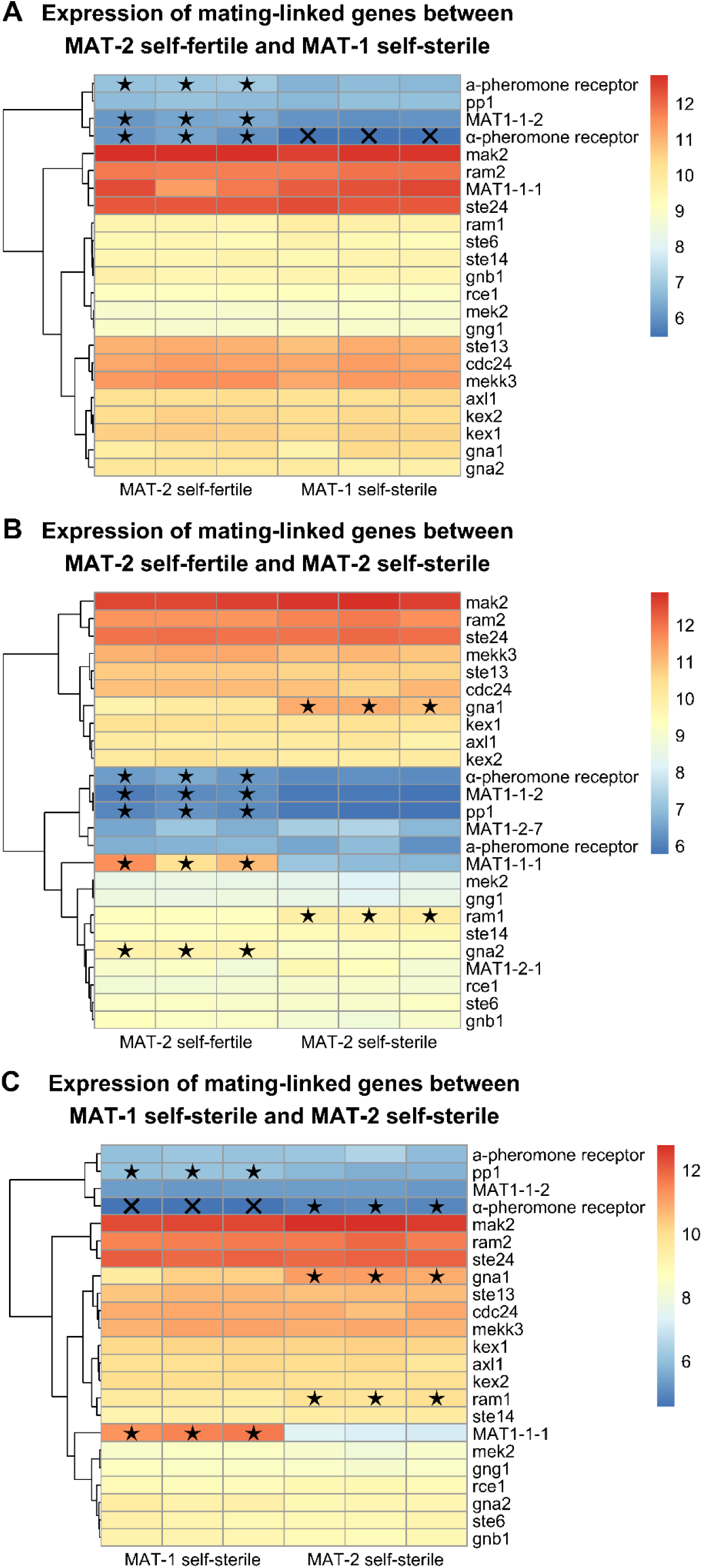
Heatmaps showing the expression of mating-related genes and genes that were significantly upregulated in an isolate compared to another (indicated by black stars). Crosses indicate genes that were not expressed in that isolate.

RNA read mapping revealed transcriptional distinctions between the *MAT1* locus configurations, offering insights into locus structure and activity. This was the case in both the self-fertile and self-sterile *MAT1* locus versions (Suppl. Fig. 3). Notably, a 5′ UTR was only detected for the *MAT1-1-1* gene in the MAT-1 self-sterile locus version. However, RNA mapped to the MAT-2 self-fertile locus configuration in the MAT-2 self-fertile and MAT-2 self-sterile isolates aligned to the *MAT1-1-1* promoter region and continued through the direct repeat and approximately 170 bp into the MAT1-2 region that is deleted during mating-type switching (Fig. 2).

**Figure 2:**
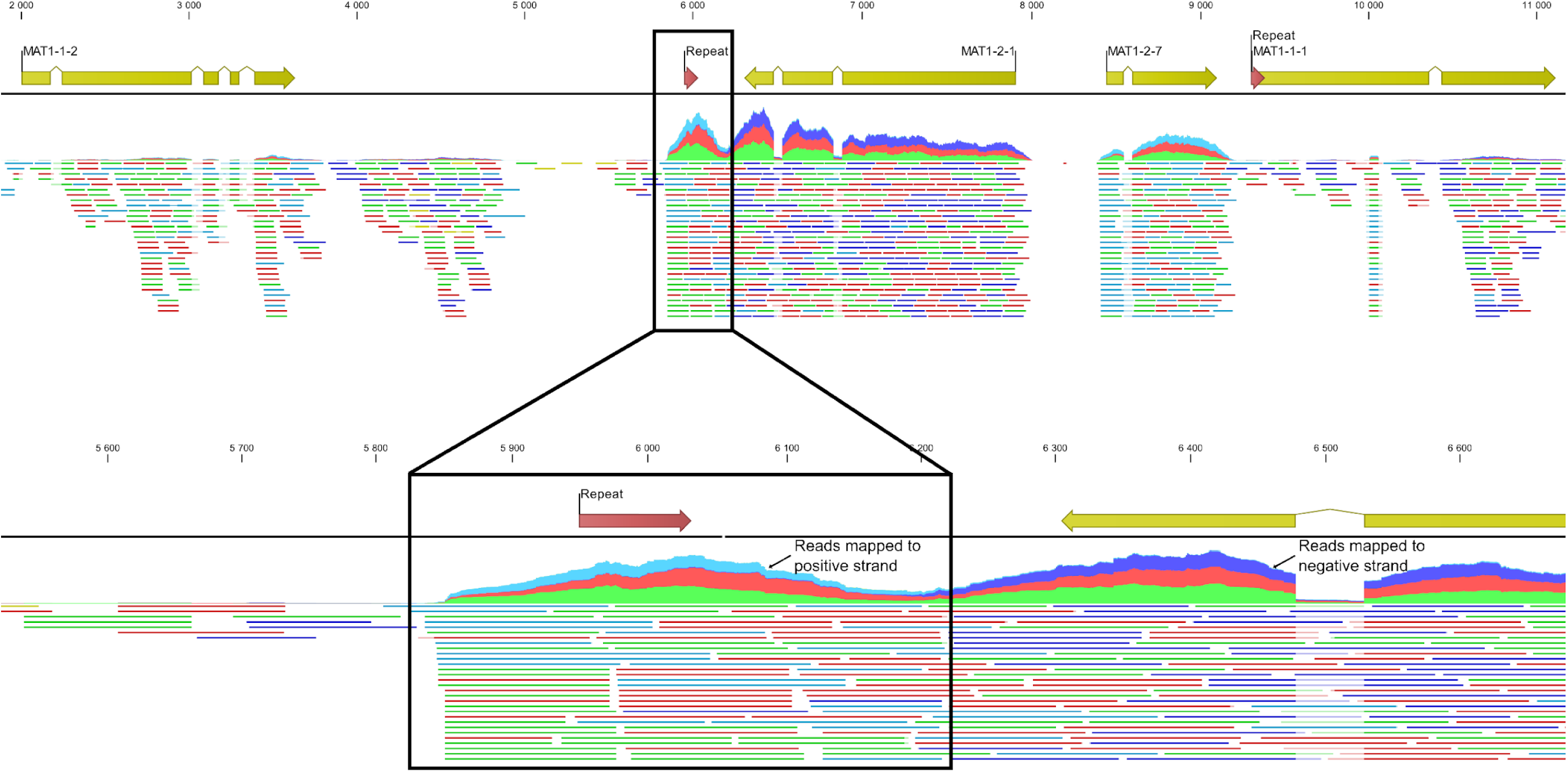
RNA reads from the MAT-2 self-sterile isolate mapped to the MAT-2 self-fertile *MAT1* locus configuration. Closer examination of the disconnected *MAT1-1-1* promotor region indicates transcripts that include this region as well as the direct repeat and some of the *MAT1-2* region. Yellow and red arrows indicate coding sequences and direct repeat sequences respectively. Light blue regions on the graph indicate reads mapping to the positive strand while dark blue regions in the graph show negative strand mapping.

#### Comparison of MAT-2 self-fertile and MAT-1 self-sterile isolates

When comparing the expression data for the MAT-1 self-sterile to the MAT-2 self-fertile isolate, 371 genes were differentially expressed. Of these, 110 genes were upregulated in the self-sterile isolate and 261 genes were upregulated in the self-fertile isolate. The top ten highly upregulated genes in the self-sterile dataset included genes involved in acetyl-CoA catabolism and metabolism, as well as genes associated with transposable elements and viruses (Fig. 3A; Suppl. Table 5). The majority of these upregulated genes in the self-fertile isolate had no detectable conserved domain, and were treated as hypothetical genes based on BLAST searches of the NCBI database (Suppl. Table 5).

**Figure 3:**
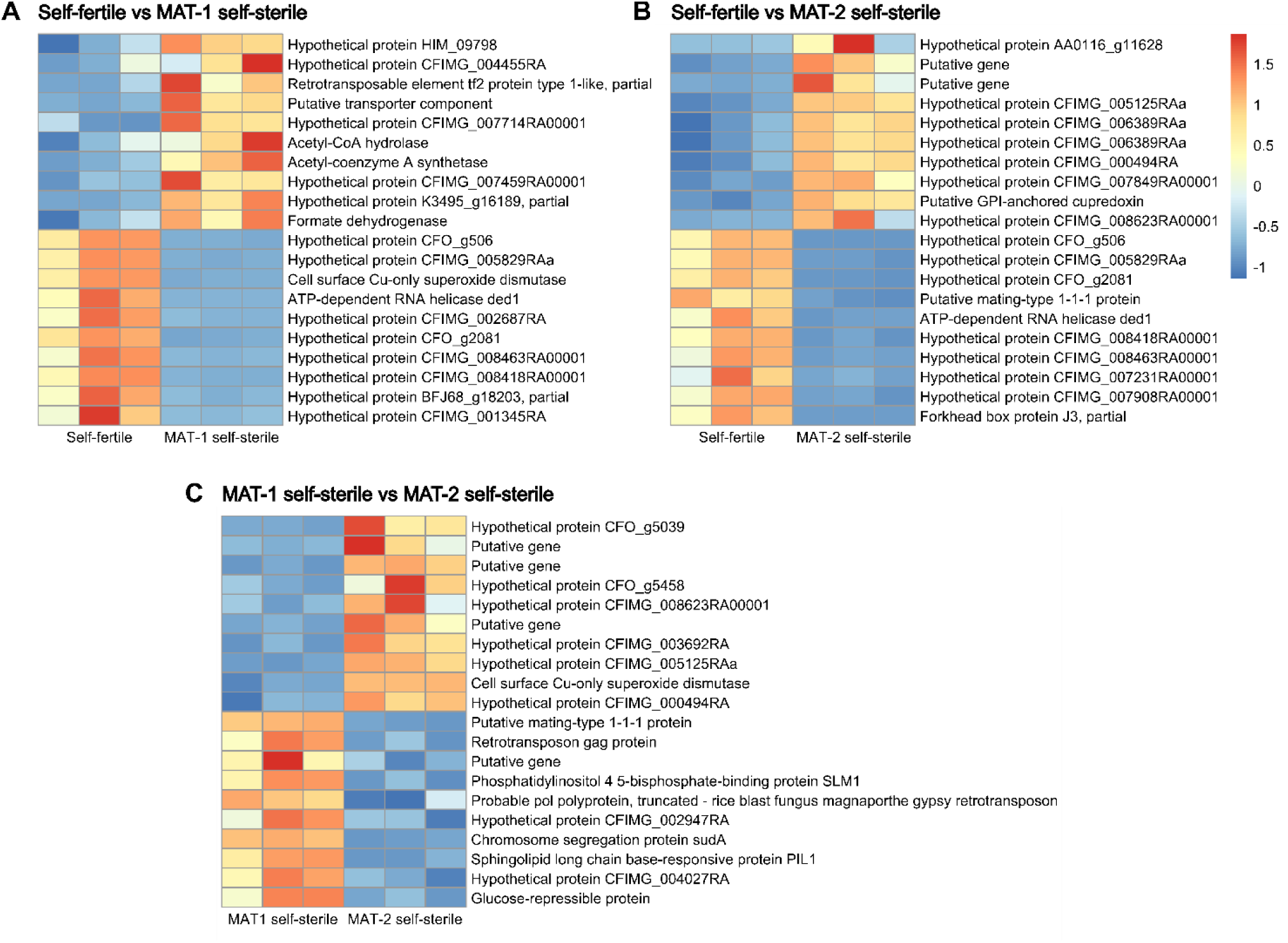
The top ten differentially expressed genes in each isolate that was compared. This is based on the log_2_ fold change where the adjusted p-value was more than 0.05. Genes labelled “putative gene” were not assigned a description during the Blast2GO pipeline.

Genes associated with the sexual cycle displayed variable expression patterns. The *MAT1-1-1* gene was expressed at similar levels in both the MAT-2 self-fertile and MAT-1 self-sterile isolates (Fig. 1A). The α-pheromone receptor was expressed only in the self-fertile isolate, whereas *MAT1-1-2* and the a-pheromone-receptor were expressed in both, but were upregulated in the self-fertile isolate. All pheromone processing genes and genes involved in the signal transduction pathway were expressed in both isolates but no significant differential expression was noted.

Twenty-six gene sets were significantly enriched in the MAT-1 self-sterile isolate compared to the MAT-2 self-fertile (Fig. 4A). These gene sets were mostly related to the production and functioning of mitochondria. These also included gene sets involved in protein synthesis and movement, as well as antibiotic metabolism. No gene sets were significantly enriched in the self-fertile isolate.

**Figure 4:**
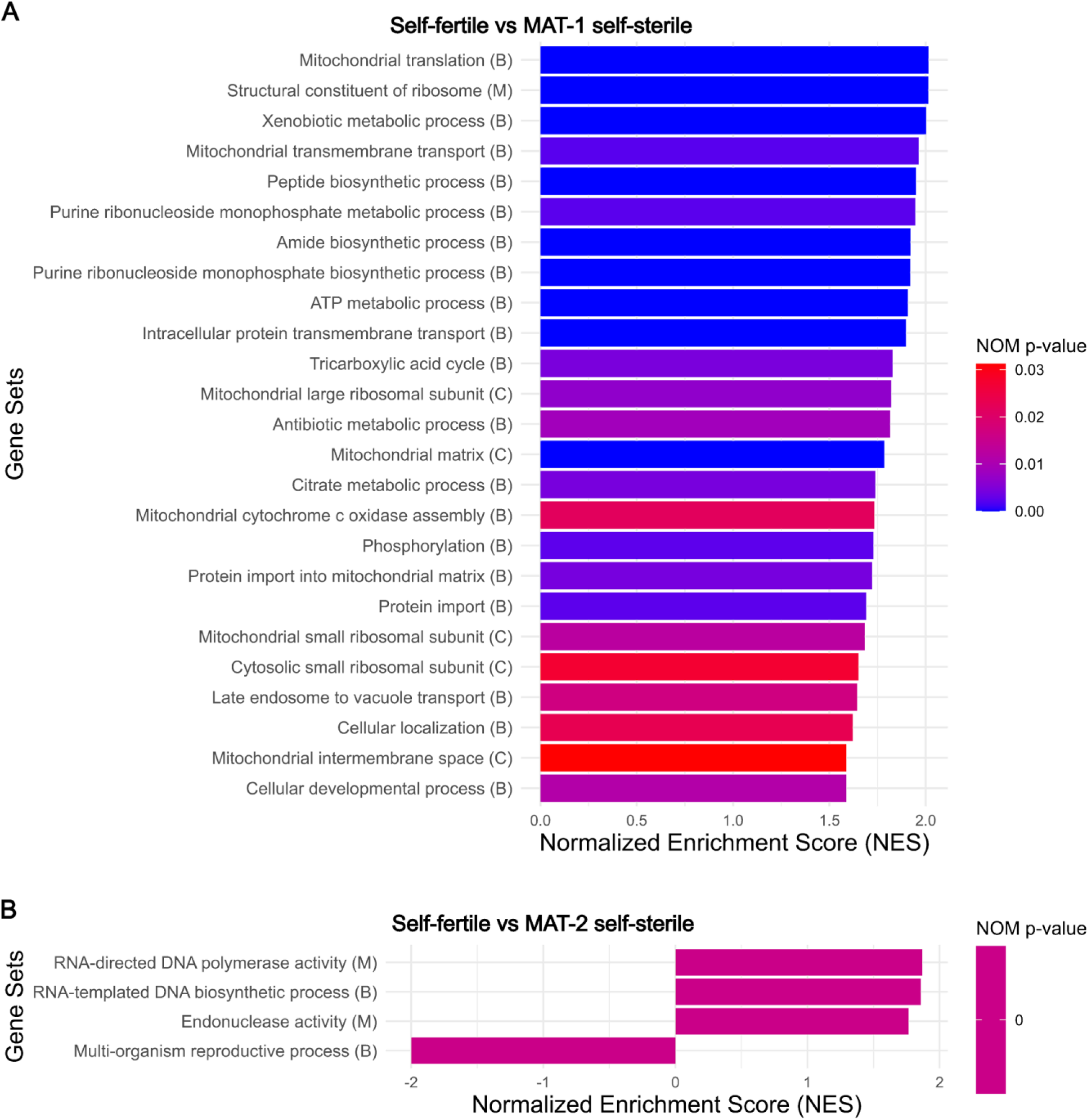
All gene sets that show significant enrichment. Letters in brackets indicate the type of GO term (B = biological process, C = cellular component, M = molecular function). (**A**) Results from preranked GSEA for the self-fertile isolate when compared the MAT-1 self-sterile isolate and (**B**) the MAT-2 self-sterile isolate.

#### Comparison of MAT-2 self-fertile and MAT-2 self-sterile isolates

A total of 1,034 genes were differentially expressed between the MAT-2 self-fertile and MAT-2 self-sterile isolates. In this comparison, more genes showed higher expression in the self-fertile isolate (573) than in the self-sterile isolate (461). The most strongly upregulated genes in the self-sterile isolate were largely annotated as hypothetical proteins with no conserved domains, with one exception containing a chromatin organisation modifier domain (Fig. 3B; Suppl. Table 5). Among the top ten upregulated genes in the self-fertile isolate were two transcription factors - *MAT1-1-1* and a forkhead box protein, while the rest were hypothetical genes, several lacking identifiable domains.

Genes linked to mating showed varied expression patterns between the two isolates. Both *MAT1-2* genes and the a-pheromone receptor were expressed at similar levels (Fig. 1B). In contrast, both *MAT1-1* genes and the α-pheromone receptor were significantly upregulated in the self-fertile isolate, with *MAT1-1-1* showing a 96-fold increase in expression relative to the self-sterile. Among pheromone processing genes, only *ram1*, which modifies the a-pheromone, was differentially expressed, with higher expression in the self-sterile isolate (Fig. 1B). Within the signal transduction pathway, *pp1* and *gnb1* were upregulated in the self-fertile isolate, while *gna1* was upregulated in the self-sterile; all other pathway components showed no significant differential expression.

Gene set enrichment analysis identified four significantly enriched gene sets in this comparison (Fig. 4B). Three of these were enriched in the self-sterile isolate and included functions related to RNA-to-DNA biosynthesis and endonuclease activity. The only gene set enriched in the self-fertile isolate was associated with reproductive processes in multicellular organisms.

#### Comparison of MAT-1 self-sterile and MAT-2 self-sterile isolates

Differential expression analysis between the MAT-1 and MAT-2 self-sterile isolates revealed 584 genes with significant changes in expression. Of these, 261 genes were upregulated in the MAT-1 self-sterile isolate, while 323 were upregulated in the MAT-2 self-sterile. Among the most highly expressed genes in the MAT-1 self-sterile were those associated with mating, transposable elements, and membrane or signalling functions (Fig. 3). In contrast, the top ten most upregulated genes in the MAT-2 self-sterile isolate were predominantly hypothetical or putative genes, lacking conserved domains and yielding no significant BLAST hits.

Although both isolates were self-sterile, key differences were observed in the expression of genes linked to the sexual cycle. The *MAT1-1-2* and a-pheromone-receptor genes were expressed at comparable levels in both isolates, whereas the α-pheromone receptor was exclusively expressed in the MAT-2 self-sterile isolate (Fig. 1C). The *MAT1-1-1* gene, expressed in both, exhibited a 140-fold higher expression in the MAT-1 self-sterile isolate. Most pheromone processing genes showed no significant differences in expression, except for *ram1*, which was upregulated in the MAT-2 self-sterile. A similar situation was seen in signal transduction pathway genes: *pp1* was more highly expressed in the MAT-1 self-sterile isolate, while *gna1* was upregulated in the MAT-2 self-sterile. No significantly enriched gene sets were identified in this comparison.

## Discussion

Mating-type genes are known as key regulators of the sexual cycle (Dyer et al. 2016; Wilson et al. 2021), but functional and chromatin immunoprecipitation studies highlight their broader roles in shaping fungal biology and development (Becker et al. 2015; Böhm et al. 2013; Doughan and Rollins 2016). In *Ceratocystis* species, unidirectional mating-type switching not only involves deletion of two *MAT1-2* genes but also appears to rearrange the locus, potentially affecting the expression of the *MAT1-1-1* gene (Wilken et al. 2024). While these genomic changes are known to directly change the sexual phenotype (Wilken et al. 2014; Yun et al. 2017), their downstream effects on gene regulation at a transcriptome-wide level, particularly in relation to self-fertile and self-sterile isolate types, remain poorly understood. In this study, we showed distinct transcriptomic signatures between three *C. albifundus* isolate types that were genetically identical, differing only at the *MAT1* locus. These included a MAT-1 self-sterile type (*MAT1-1* genes only), a MAT-2 self-sterile type (the literature-defined “MAT-2 self-fertile” locus with *MAT1-1* and *MAT1-2* genes), and a MAT-2 self-fertile heterokaryon containing both locus versions seen in the self-sterile isolates. There was clear differential expression of the *MAT* genes and associated components of the sexual signalling pathway, including pheromone precursors and their receptors. These findings underscore the central role of mating-type genes and their downstream networks in mediating broader transcriptional responses. They also suggest that unidirectional mating-type switching may have biological consequences that extend well beyond the regulation of sexual reproduction.

Previous research has proposed that unidirectional mating-type switching joins *MAT1-1-1* with its promoter, potentially affecting its expression profile (Wilken et al. 2024). In the self-fertile configuration, *MAT1-2* genes lie between *MAT1-1-1* and its promoter, blocking transcription. Switching removes this intervening region, restoring promoter proximity and enabling *MAT1-1-1* expression. This mechanism is similar to that in other fungi where insertion of *MAT1-2* disrupts the *MAT1-1-1* coding sequence (Krämer et al. 2021; Xu et al. 2016; Yun et al. 2017). In our RNA-seq data, *MAT1-1-1* was expressed at similar levels in the MAT-2 self-fertile and MAT-1 self-sterile isolates, both of which possess the promoter-linked version of the gene. These results support the model that switching enables *MAT1-1-1* transcription (Wilken et al. 2024), facilitating the coexistence of nuclei expressing both core mating-type genes in a single individual. Since *MAT1-1-1* and *MAT1-2-1* are primarily responsible for sexual reproduction (Wilson et al. 2021), switching may serve as a mechanism to ensure their concurrent expression within a single individual.

Interestingly, partial transcripts from *MAT1-1-1* were detected in the MAT-2 self-sterile isolate (Fig. 2), despite this isolate carrying the self-fertile *MAT1* locus configuration (Engelbrecht and Harrington 2017; Wilken 2015). Transcript mapping showed high read density at the *MAT1-1-1* promoter but little beyond it, suggesting that transcription is initiated but prematurely terminated due to it continuing into intergenic sequence rather than *MAT1-1-1*. These results imply that while *MAT1-1-1* is constitutively induced, a functional transcript is only produced when it is connected to its promoter, as in the MAT-1 self-sterile locus. However, a small number of reads did map to the *MAT1-1-1* coding region, and the gene was detected as expressed. This could possibly reflect a low-level of successful switching, a view supported by minimal PCR amplification of the MAT-1 self-sterile locus in these isolates. Although a *MAT1* locus version allowing *MAT1-1-1* gene expression was therefore present, these isolates remained phenotypically self-sterile. Collectively, these results suggest that sexual reproduction in switching species may not depend only on the presence of both locus configurations, but also on their relative abundance within the mycelium.

Elevated expression of *MAT1-1-2* in self-fertile isolates point to a role beyond mating-type determination, potentially in later stages of sexual development. In this study, *MAT1-1-2* was expressed in all isolates but was notably upregulated in the MAT-2 self-fertile isolate. While its presence in self-sterile isolates could reflect a function in mating-type identity, increased expression in the self-fertile isolate, where sexual development is active, suggests a role beyond mate recognition. Secondary *MAT* genes like *MAT1-1-2* have been associated with post-fertilization processes, such as fruiting body development (Arnaise et al. 2001; Rodenburg et al. 2018; Wilson et al. 2020). Uniquely, *MAT1-1-2* is the only *MAT* gene retained with its flanking regions in both self-fertile and self-sterile configurations across *Ceratocystidaceae* species capable of switching (Wilken et al. 2014). Its conserved genomic position and expression profile make the *MAT1-1-2* gene a strong candidate for functional studies aimed at understanding its role in the sexual cycle.

Pheromone receptor expression in *C. albifundus* appears to be mating-type dependent and linked to sexual activity. While these receptors are typically expressed regardless of mating type (Kim and Borkovich 2004; Kim et al. 2012), some fungi do show mating-type specific expression, where *MAT1-1* and *MAT1-2* genes control a- and α-pheromone receptor expression, respectively (Shen et al. 1999; Wilson et al. 2020; Zhang et al. 1998). In our study, the α-pheromone receptor was absent in the MAT-1 self-sterile isolate, likely due to the loss of *MAT1-2* genes through mating-type switching. This supports the idea that α-pheromone receptor expression is dependent on *MAT1-2* genes. Both receptor genes were expressed in the MAT-2 self-fertile isolate and were generally upregulated relative to the self-sterile isolates, possibly reflecting an active role in the sexual cycle. These patterns parallel those seen in *Huntiella moniliformis*, where α-pheromone receptor expression increases during active sexual reproduction (Wilson et al. 2018).

Pheromone gene expression did not match that of their cognate receptors, a discrepancy likely influenced by the timing of RNA sampling. Neither the a- nor α-pheromone genes were expressed in the MAT-2 self-fertile nor in either of the self-sterile isolates at the sampled time point. Although pheromone expression does not follow a strict temporal pattern, it is generally observed prior to or during sexual reproduction (Coppin et al. 2005; Kim and Borkovich 2006; Pöggeler 2000). In this study, a single time point was used to compare transcriptomic differences, with self-sterile isolates grown alone showing no signs of sexual development, while the self-fertile isolate was actively producing ascomata. Previous studies have shown that sexual reproduction can occur when MAT-1 and MAT-2 self-sterile isolates are paired (Engelbrecht and Harrington 2017; Harrington and McNew 1997), suggesting that pheromone expression may be triggered by interaction between compatible partners. The lack of expression in isolated self-sterile cultures highlights the need for time-course studies of mating interactions to better understand pheromone dynamics in switching-capable species.

While this study focused on mating-related gene expression, most differentially expressed genes were not directly linked to mate recognition or reproduction. This was reflected in gene set enrichment results, where only the “multi-organism reproductive process” category was enriched in the MAT-2 self-fertile isolate compared to its MAT-2 self-sterile counterpart. Given that the isolates are genetically identical apart from the *MAT1* locus, the widespread transcriptomic changes observed are striking. Many of the upregulated genes in the self-fertile isolate may contribute to sexual reproduction indirectly (Coppin et al. 1997; Wilson et al. 2019), as this phase involves extensive morphological, metabolic, and chemical changes that also occur during other stages of the fungal life cycle. Notably, the two self-sterile, vegetative isolates also differed substantially in gene expression, suggesting that the *MAT1* locus influences a wider range of biological processes. Previous studies have linked this region to traits such as conidial formation, growth, hyphal morphology, pathogenicity, spore dimorphism (Böhm et al. 2013; Lee et al. 2015; Lockhart et al. 2005; Njambere et al. 2011; Zhan et al. 2007), emphasising the need for functional studies of mating-type genes and locus architecture in pathogenic fungi like *C. albifundus*.

## Data availability statement

RNA generated in this study can be accessed on NCBI’s Sequence Read Archive.

## Acknowledgements

We thank Dr Magriet van der Nest and Ms Vinolia Danke for the access to the annotated genome used in this study. Portions of this manuscript’s text were refined with the assistance of ChatGPT (OpenAI), which was used to improve clarity, conciseness, and language flow. No content, data interpretation, or scientific conclusions were generated by the tool. The authors take full responsibility for the manuscript’s content and accuracy.

## Supplementary Tables

**Supplementary table 1:**
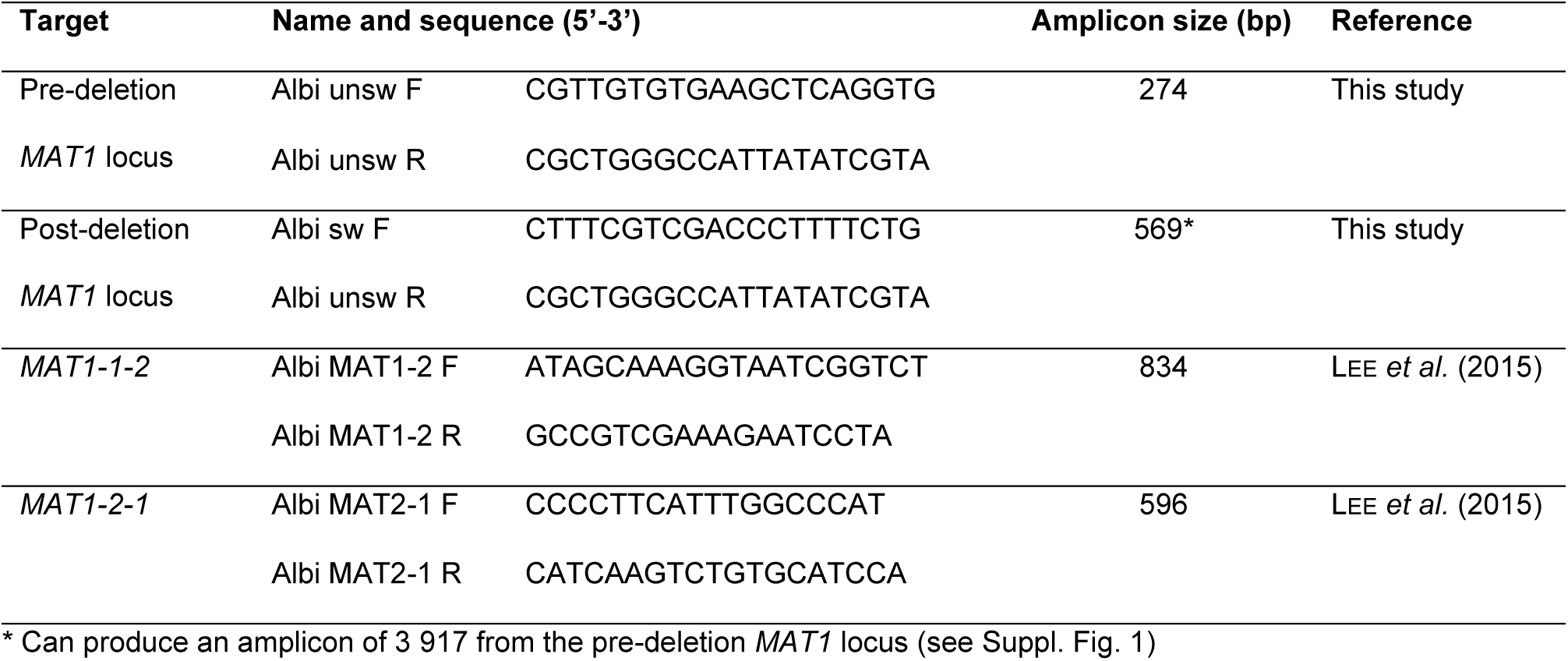
The primer sets used to screen for MAT-1 and MAT-2 self-sterile isolates.

**Supplementary table 2:** CLC Genomics default settings for “Trim Reads” function

|  |  |
| --- | --- |
| **Quality trimming** |  |
| Trim using quality scores | Yes, quality limit: 0.05 |
| Trim ambiguous nucleotides | Yes, maximum number of ambiguities: 2 |
| **Adaptor trimming** |  |
| Automatic read-through adaptor trimming | Yes |
| **Homopolymer trimming** |  |
| Trim homopolymers from 5' | No |
| Trim homopolymers from 3' | No |
| **Sequence filtering** |  |
| Remove fixed number of bases from 5' terminal | No |
| Remove fixed number of bases from 3' terminal | No |
| Remove bases until reads are a fixed length | No |
| Filter on length | No |

**Supplementary table 3:**
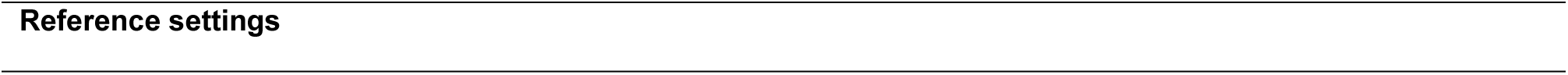
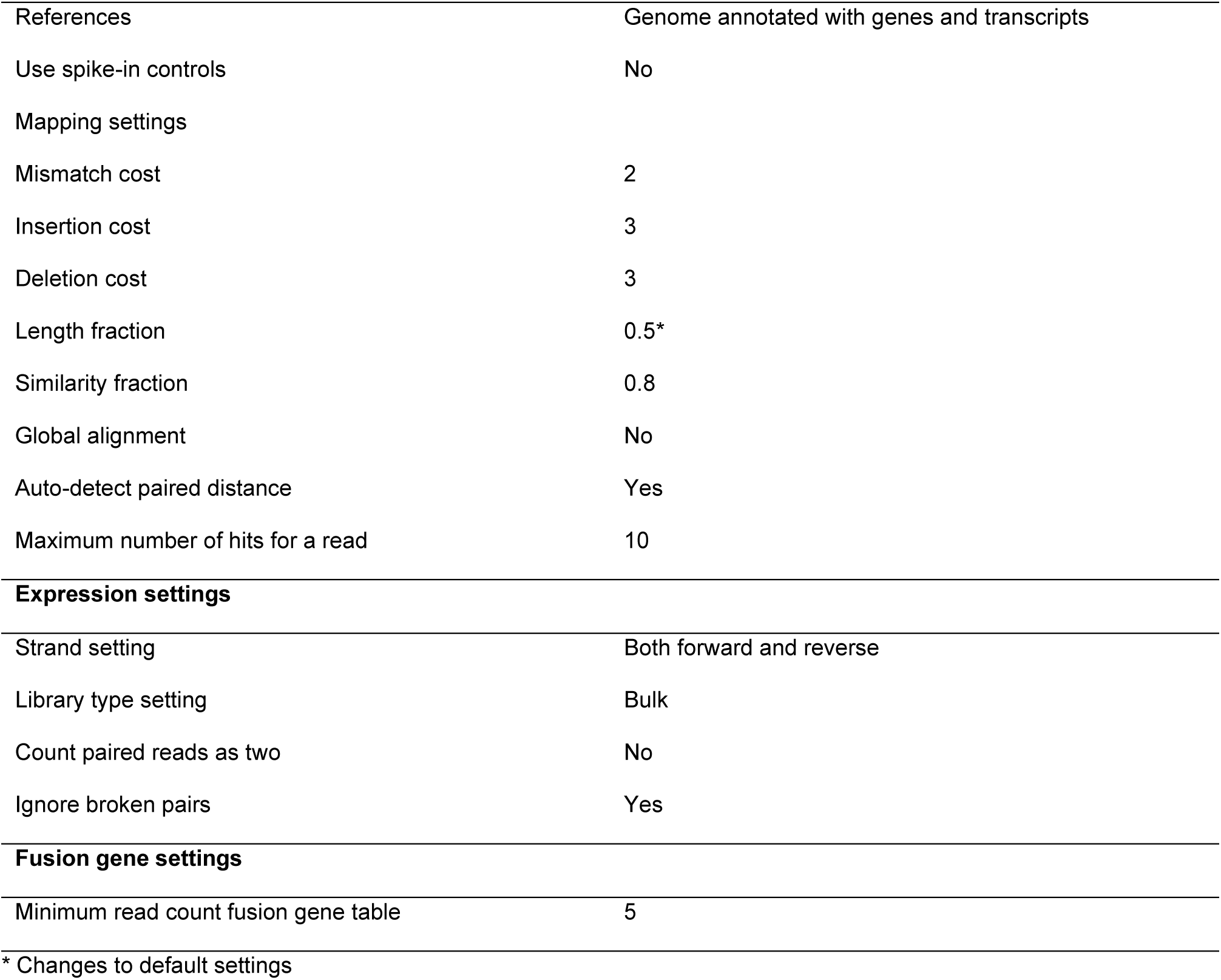
CLC Genomics settings for “RNA-Seq Analysis” function used

|  |
| --- |
| **Reference settings** |

|  |  |
| --- | --- |
| References | Genome annotated with genes and transcripts |
| Use spike-in controls | No |
| Mapping settings |  |
| Mismatch cost | 2 |
| Insertion cost | 3 |
| Deletion cost | 3 |
| Length fraction | 0.5* |
| Similarity fraction | 0.8 |
| Global alignment | No |
| Auto-detect paired distance | Yes |
| Maximum number of hits for a read | 10 |
| **Expression settings** |  |
| Strand setting | Both forward and reverse |
| Library type setting | Bulk |
| Count paired reads as two | No |
| Ignore broken pairs | Yes |
| **Fusion gene settings** |  |
| Minimum read count fusion gene table | 5 |

\* Changes to default settings

**Supplementary table 4:** Details on RNA-seq data

| Samples | Concentration<br>(ng/ul) | Raw reads<br>(mil) | Trimmed<br>reads (mil) | Reads mapped in pairs<br>and broken pairs (%) | Transcripts<br>(mil) |
| --- | --- | --- | --- | --- | --- |
| MAT-2 self-fertile rep 1 | 188.3 | 48.9 | 48.9 | 87.90 | 10.5 |
| MAT-2 self-fertile rep 2 | 435.5 | 55.1 | 55.1 | 89.93 | 12.1 |
| MAT-2 self-fertile rep 3 | 765.0 | 37.4 | 37.4 | 87.92 | 8.0 |
| MAT-1 self-sterile rep 1 | 681.9 | 43.6 | 43.6 | 90.11 | 9.6 |
| MAT-1 self-sterile rep 2 | 976.3 | 44.2 | 44.2 | 90.75 | 10.0 |
| MAT-1 self-sterile rep 3 | 846.8 | 46.3 | 46.3 | 85.50 | 9.2 |
| MAT-2 self-sterile rep 1 | 659.4 | 33.6 | 33.6 | 91.07 | 7.1 |
| MAT-2 self-sterile rep 2 | 790.2 | 35.5 | 33.5 | 90.62 | 7.2 |
| MAT-2 self-sterile rep 3 | 1 489.0 | 37.7 | 37.7 | 89.60 | 8.0 |

**Supplementary table 5:**
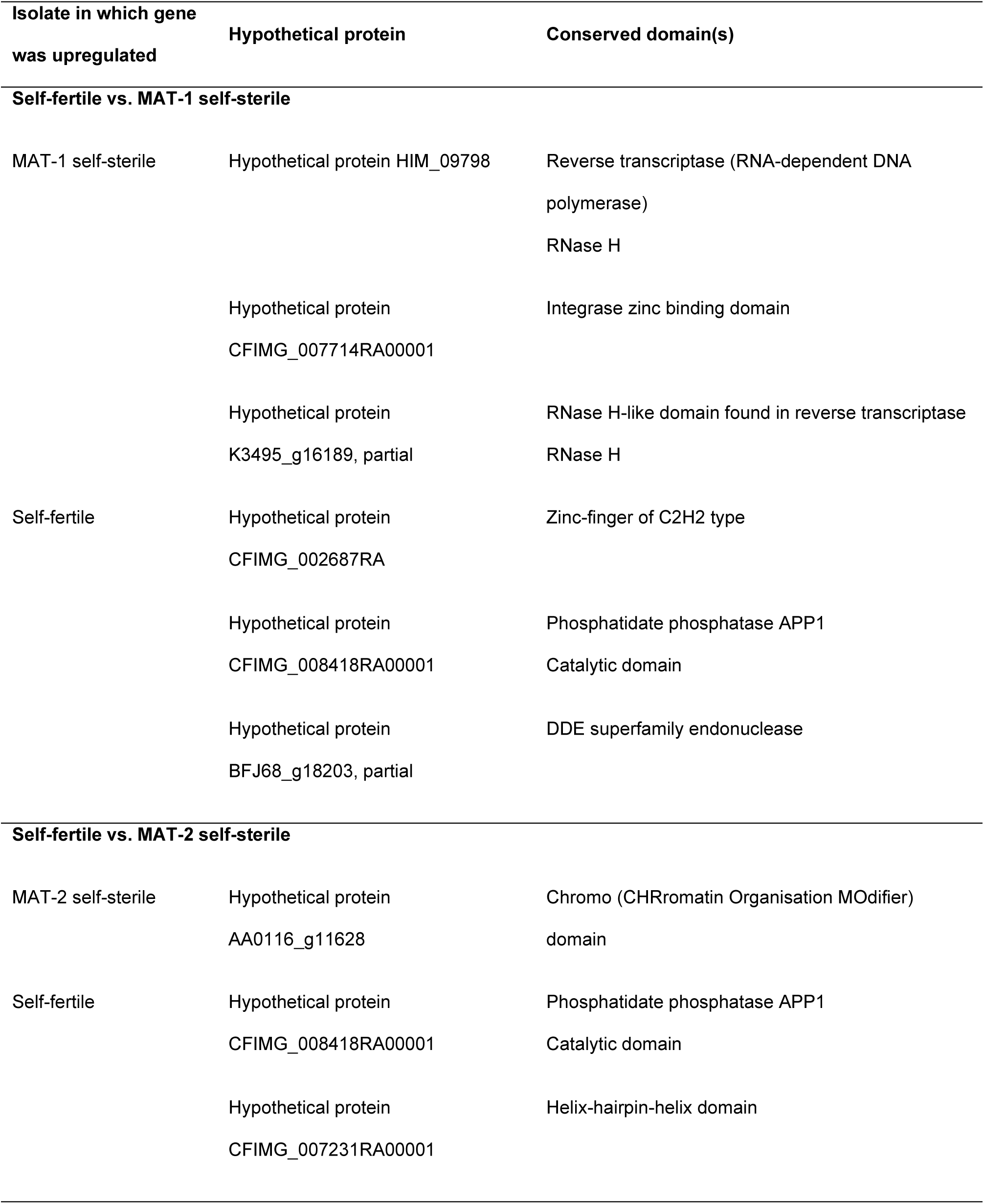
Hypothetical genes that were part of the top ten differentially expressed genes for each isolate in each comparison for which a conserved domain could be identified

## Supplementary Figures

**Supplementary Figure 1:**
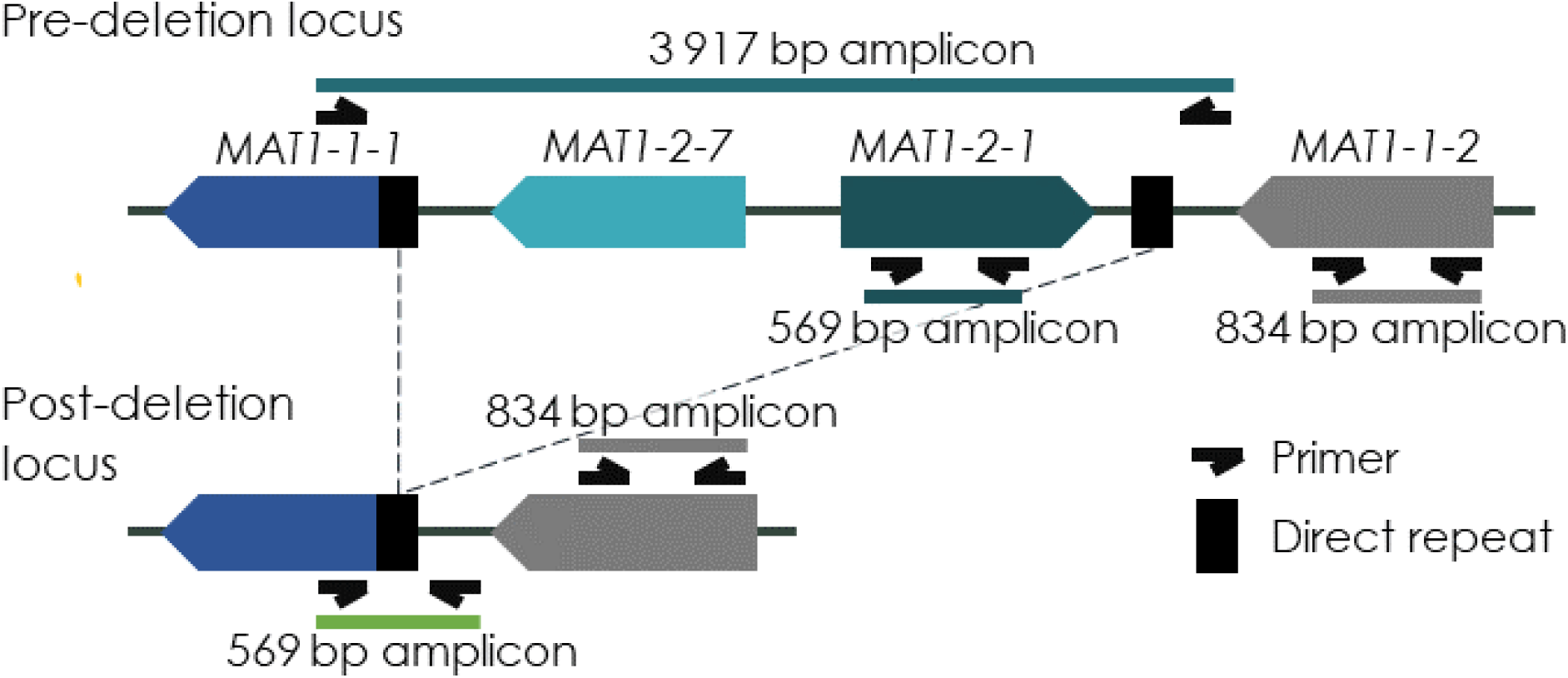
A visual representation of the two versions of *MAT1* locus with the binding sites and expected amplicons from the primer sets used to determine the fertility type of an isolate. This figure is not drawn to scale. Primers used can be found in Suppl. Table 1.

**Supplementary Figure 2:**
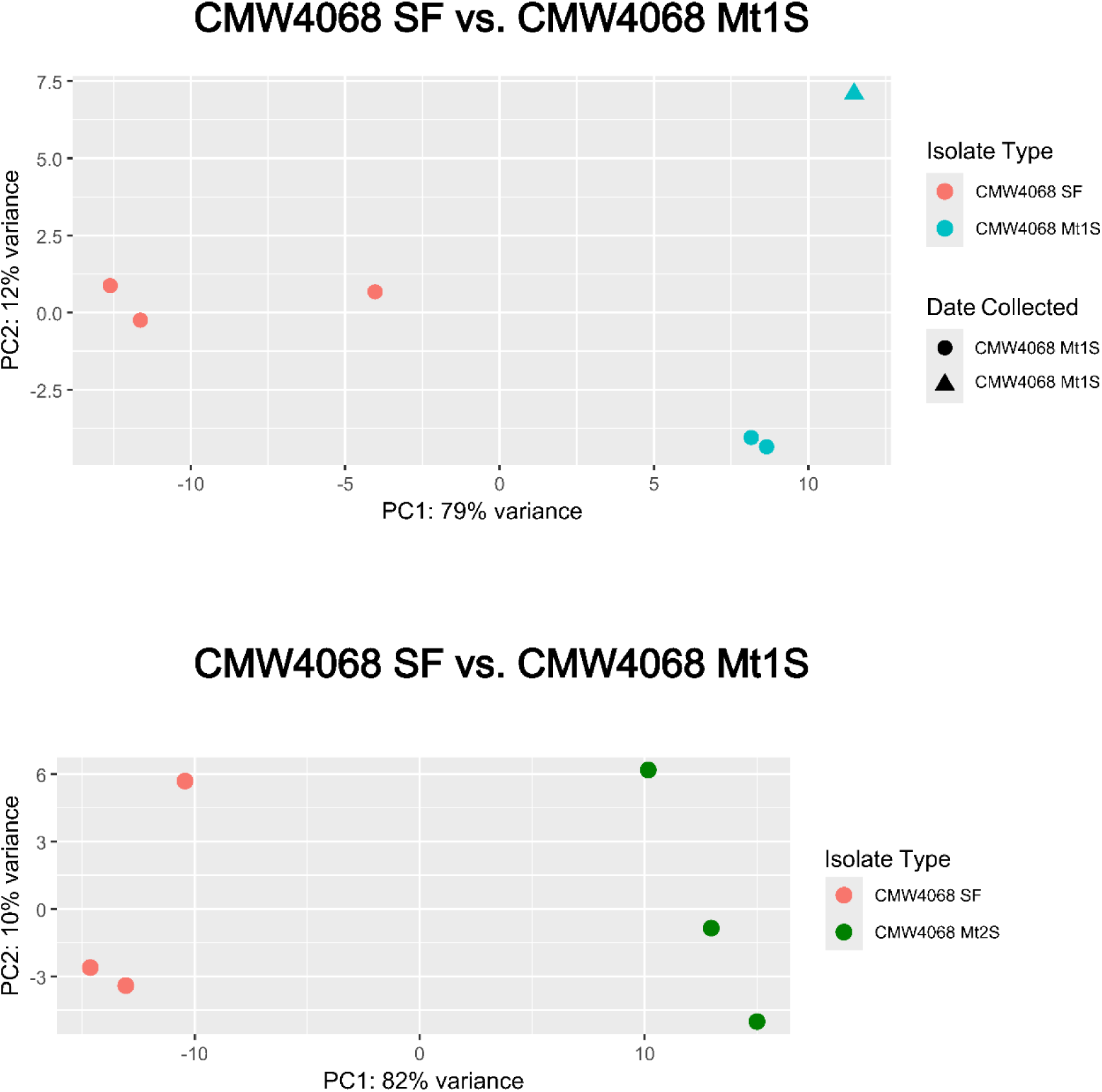
PCA plots of regularised-logarithmic (rlog) transformation data sets.

**Supplementary Figure 3:**
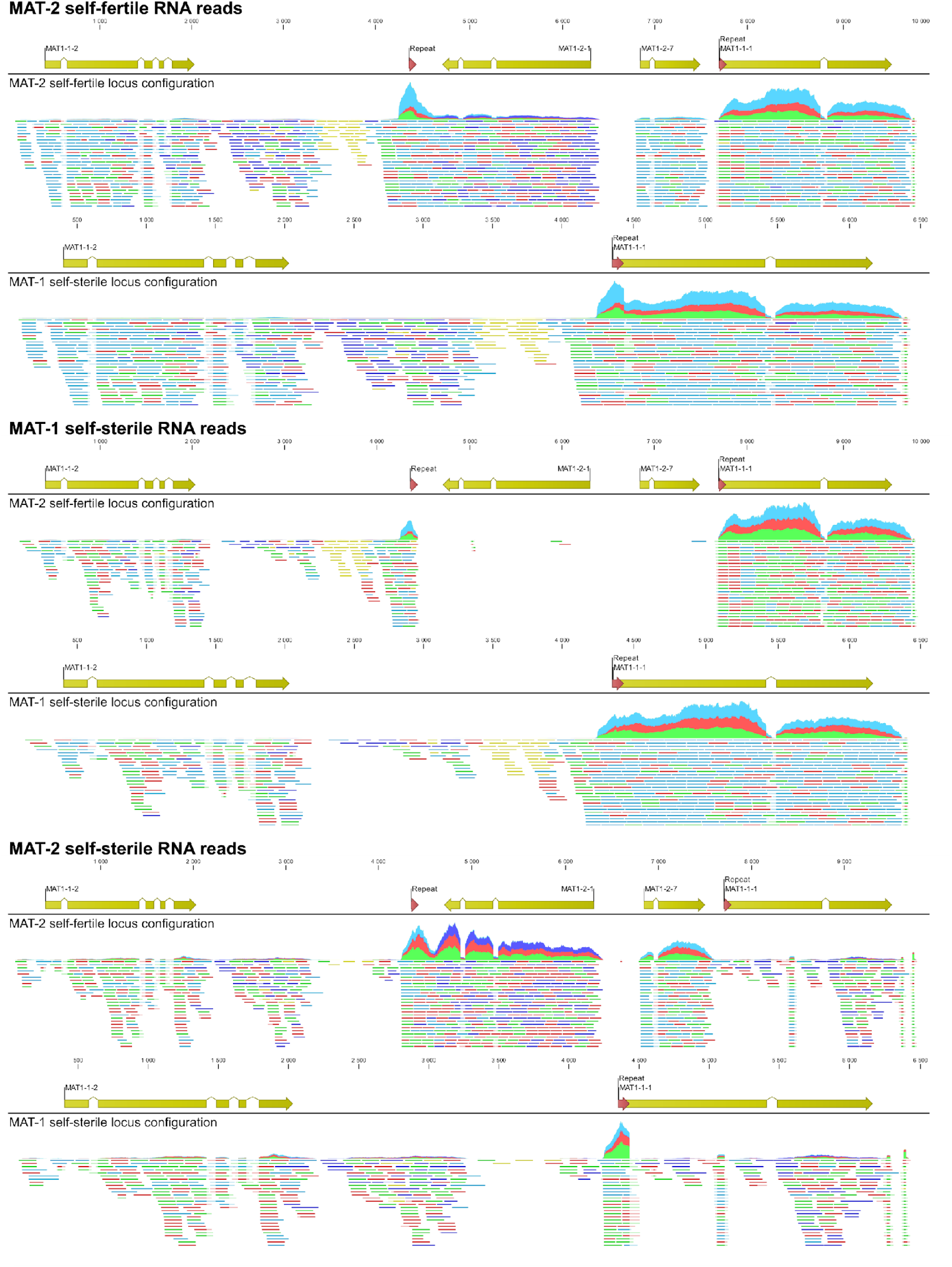
RNA reads mapped to the MAT-2 self-fertile and MAT-1 self-sterile mating-type locus configuration. Yellow arrows indicate genes and red arrows indicate direct repeat regions.

## Supplementary Files

**Supplementary File 1:** Methods used to produce various plots and heatmaps

Dispersion trends were determined through regularised-logarithmic transformation (rlog function on DESeq2), the results of which were used for principal component analyses (plotPCA function). Heatmaps of the top 10 differentially expressed genes that had the largest log_2_ fold change in each isolate were made using pheatmap (v. 1.0.12). The heatmaps were based on the rlog-transformed normalised counts.

**Supplementary File 2:** Reason for variation between technical replicates of CMW4068 Mt1S

With the exception of a single sample of CMW4068 Mt1S, all RNA isolations were performed on the same day. The one CMW4068 Mt1S RNA isolation was performed at a later date due loss of a single sample during the original isolations. The experimental design was performed in the same way as all other samples, however, the variation we see between CMW4068 Mt1S technical replicates (Suppl. Fig. 3) was likely due to batch effect.

